# Glutamatergic Dysfunction of Astrocytes in Paraventricular Nucleus of Thalamus Contributes to Adult Anxiety Susceptibility in Adolescent Ethanol Exposed Mice

**DOI:** 10.1101/2025.05.20.654912

**Authors:** Aubrey Bennett, Hyunjung Kim, David Thomas, Peter Biggs, Roxan Ara, Asamoah Bosomtwi, Seungwoo Kang

## Abstract

Repeated ethanol exposure during adolescence increases the risk for displaying anxiogenic phenotype in adulthood, but the underlying mechanisms are not fully understood. The paraventricular nucleus of thalamus (PVT) has been considered a hub brain area for controlling the anxiety network in the brain. Recent structural and functional investigations indicate that the PVT exhibits diverse neural signals aligned with early-life events, which are highly linked with anxiety-like behaviors. However, it remains unknown if repeated ethanol exposure during adolescence will affect the coordinated brain activities of the PVT in adulthood, and consequent behavioral adaptation. Here we show that adolescent repeated intermittent ethanol exposure (AIE) triggers anxiety-like behaviors and parallelly induces the glutamatergic adaptation in the PVT after four weeks withdrawal from the last ethanol exposure. The firing rates, along with the spatiotemporal calcium transients in the PVT neurons during behavior were increased in the AIE mice compared to those in the counterpart control mice. Importantly, with the chemogenetic inhibition of the PVT neurons, we found alleviated the anxiety-like behavior in the AIE mice.

The increased neuronal activities in the PVT of AIE mice was, at least partly, via the reduction of GLT1 (an astrocyte dominant glutamate transporter, known as EAAT2, slc1a2). Our non-invasive magnetic resonance spectroscopy (MRS) measures also showed an increase in glutamate/GABA ratio in the thalamic area including the PVT of the GLT1 conditional knock-down mice, which exhibited the heightened anxiety-like behavior. In addition, while the selective knock-out of GLT1 in the astrocytes of PVT in the alcohol naïve mice induces anxiogenic phenotypes, the selective upregulation of GLT1 in the PVT astrocytes of the mice that were treated with AIE paradigm alleviated the anxiety-like behaviors as well. These findings highlight the significant role of PVT astrocytic GLT1 in the anxiogenic phenotype in adulthood induced by withdrawal from adolescent repeated ethanol exposure, suggesting that GLT1 in the PVT could serve as a therapeutic target for alcohol use disorder and comorbid emotional disorders.

## Introduction

Adolescence is a crucial period for the maturation of brain [1–4]. As a result, those who are exposed to alcohol at this early period are more likely to experience many psychological and physical issues even into adulthood [5–13]. One remarkable consequence of exposure to alcohol on the adolescent brain is the fact that it increases the likelihood developing generalized anxiety disorder later in life [14–18], which in adults is also associated with alcohol use disorder (AUD) [19]. AUD and anxiety are indeed not only common comorbidities but also contribute to each other’s development [13,19]. However, the neurobiological mechanisms underlying adolescent chronic intermittent alcohol exposure-induced adult behavioral susceptibility in anxiety-like behavior remain unclear.

The thalamic circuits mediating arousal have been recognized as a link between the adolescent experience and the molecular and behavioral traits associated with psychiatric disorders in adults [20–23]. Especially, the paraventricular nucleus of thalamus (PVT) has been considered a hub of anxiety network [24,25] because anxiety is characterized as a state of alertness and hypervigilance to external environmental cues [26]. Indeed, adverse experiences in early life induce the pathophysiological adaptation in the PVT, and consequent anxiogenic profiles in rodent models [23,27]. However, it remains unknown if the repeated ethanol exposure during adolescence will affect the coordinated brain activities of the PVT in adulthood, and consequent behavioral adaptation.

In the current study, using chemogenetics, electrophysiology, calcium imaging, magnetic resonance spectroscopy, and behavioral evaluation approaches with transgenic mouse models, we revealed how the AIE leads an adaptation in glutamatergic signatures via the astrocyte-neuron interaction in the PVT and a subsequent behavioral adaptation.

## Materials and Methods

### Animals

All experimental procedures received approval from the Augusta University Institutional Animal Care and Use Committee and were conducted in accordance with NIH guidelines. C57BL/6J mouse line (Catalog no. 000664) and the GFAP^cre/+^ line [Catalog no. 024098; B6.Cg-Tg(Gfap-cre)77.6Mvs/2J] were acquired from Jackson Laboratory (Bar Harbor, ME). GLT1-flox mice (Catalog no. 026619; B6.Cg-Slc1a2^tm1.1Ncd^/J) [28] were received from Drs. Doo-Sup Choi at Mayo Clinic and Niels Christian Danbolt at University of Oslo. Ai9-GLT1 mice for cre-dependent overexpression of GLT1 were received from Drs. Doo-Sup Choi at Mayo Clinic and Ho Lee at Korea Cancer Center. Mice were accommodated in standard Plexiglas cages. The colony room was regulated to a consistent temperature of 24 ± 1°C and humidity of 60 ± 2%, with a 12-hour light/dark cycle (lights on at 07:00 A.M.). Mice were provided *ad libitum* access to food and water. For all the experiments, we used both male and female mice.

### Stereotaxic surgery for virus injection

Mice were anesthetized with isoflurane (1.5% in oxygen) and positioned on the rotational digital stereotaxic apparatus (RWD Life Science). The skull was aligned with a dual-tilt position equalizer and holes were drilled in the skull at the designated stereotaxic coordinates. Viruses were infused to the posterior parts of the PVT (AP −1.6 mm, ML +0.0 mm, DV −3.0 mm from bregma) at a rate of 100 nl/min for 3 minutes via a 34-gauge needle (Catalog no. NF34BV; World Precision Instruments) with a micro-syringe pump (World Precision Instruments). The injection needle was kept in place for an extra 10 minutes following the injection. We injected the AAVs at the following titers: AAV_5_-CaMKIIa-GCaMP6s, 4.7×10^12^ GC/ml (UNC Vector Core); AAV_5_-CaMKIIa-hM4Di-mCherry, 2.4×10^13^ GC/ml (Addgene); AAV_5_-GFAP-mCherry, 3.1×10^12^ GC/ml (Vector Biolab); AAV_5_-GFAP-mCherry-Cre, 4.3×10^12^ GC/ml (UNC Vector Core). We administered buprenorphine sustained release (1.3 mg/kg, s.c.; CoVetrus) to mitigate postoperative pain.

### Brain slice preparation and *ex vivo* electrophysiology

Brain slices containing the PVT region were prepared for electrophysiological recordings, as described [29]. Briefly, mice were deeply anesthetized by isoflurane inhalation, after which the brain was rapidly extracted and immersed in ice-cold sucrose-based artificial cerebrospinal fluid (aCSF) containing the following (in mM): 87 NaCl, 75 sucrose, 2.5 KCl, 11.25 NaH_2_PO_4_, 0.5 CaCl_2_, 7 MgCl_2_, 25 NaHCO_3_, 0.3 l-ascorbate, and 25 glucose, and oxygenated with 95% O_2_/5% CO_2_. Coronal brain slices (300-350 μm) were cut with a vibrating compresstome (VF-310-0Z, Precisionary Instruments), subsequently placed in a slice holding chamber and incubated for 30 min at 34°C and kept for at least 1 hour at room temperature (24-25°C) in carbonated (95% O_2_/5% CO_2_) standard artificial cerebrospinal fluid (aCSF) containing the following (in mM): 126 NaCl, 1.25 NaH_2_PO_4_, 1 MgCl_2_, 2 CaCl_2_, 2.5 KCl, 25 NaHCO_3_, and 11 glucose. Electrical signals were captured using an Axon 700B amplifier, a Digidata 1550B A/D converter, and Clampfit 11.0 software (Molecular Devices). Throughout the experiments, the bath was consistently perfused with warm (32 °C) carbonated aCSF at a rate of 2.0-2.5 ml/min. Patch pipettes had a resistance of 4-6 MΩ when filled with a solution containing (in mM): 140 Cs-methanesulfonate, 5 KCl, 2 MgCl_2_, 10 HEPES, 2 MgATP and 0.2 Na_2_GTP for voltage clamping. The pH was adjusted to 7.2 with Tris-base and the osmolality to 310 mOsmol/L with sucrose. Healthy cells were identified using a high magnification microscope (at 400X, Nikon FN1 Microscope, Melville, NY) based on their morphology (round, ovoid, and non-swelled plasma membrane). The spontaneous firing was recorded using the loose-patch cell-attached method.

### *In vivo* electrophysiology

The *in vivo* electrophyiological recordings were performed as described [29,30]. Briefly, mice were anesthetized by urethane (1.5 g/kg, *i.p.*; Sigma-Aldrich) [29] and positioned on a stereotaxic frame (RWD Life Science). We continuously monitored respiration rate and pedal withdrawal response during recordings, while the body temperature was maintained using a small-animal feedback-controlled warming pad (Kent Scientific Corporation). Small burr holes on skulls were created to insert high-impedance microelectrodes (Cambridge NeuroTech). The reference wire (Ag/AgCl, 0.03 inches in diameter, A-M systems) was positioned in the contralateral parietal cortex. Electrophysiological signals were digitized at 20 kHz and band pass–filtered from 300 to 3000 Hz (Intan Technologies). The data were analyzed with Clampfit (version 11.2, Molecular Devices) and a custom-written code in MATLAB (R2019a, The MathWorks).

### *In vivo* Ca^2+^ signal with fiber-photometry

We monitored cellular Ca^2+^ transients in real-time *in vivo* by fiber-photometry as described [15,30]. Three weeks post-injection of the AAV encoding GCaMP6s in the PVT, we implanted a fiber optic cannulae into the PVT (AP −1.40 mm, ML +0.0 mm, DV −2.8 mm from bregma). The fluorescence signals were captured at 30 frames per second using a fiber-photometry system (Plexon Multi-Fiber Photometry System, Plexon, Dallas, Texas), which had a dichroic mirror and a lens linke to a photomultiplier tube (PMT) to refect beams from 465 nm and 410 nm LED wavelengths. The collected data were analyzed using the MatLab-based photometry modular analysis program, pMAT [31].

### Chemogenetics and drug treatments

JHU31670 was purchased from Hello Bio (J60; Princeton, NJ), noted for its superior blood-brain barrier penetrance and enhanced affinity, potency, and selectivity of DREADDs (Designer Receptors Exclusively Activated by Designer Drugs) [32]. For the activation of DREADDs, we administered J60 (0.3 mg/kg) or saline intraperitoneally 10 minutes prior to the experiments in mice. These concentration have been demonstrated to have no off-target effects in rodent behavioral tests [32–36].

### Magnetic Resonance Spectroscopy (MRS)

All experiments were conducted using a Bruker Biospec 7.0 Tesla 30 cm horizontal bore scanner (Bruker Biospin MRI GmbH, Germany), featuring a BGA12SHP gradient system that generates pulse gradients of 660 mT/m along each of the three axes, equipped with AVANCE III HD electronics and interfaced to a Bruker Paravision 6.0.1 console. A 4-channel surface phase array coil (Rapid MRI International, Columbus OH) served as the receiver, while a Bruker 86 mm linear-volume coil as the transmitter. Throughout the recordings, the animal was maintained under 1–2% isoflurane anesthesia and respiratory rate and body temperature were consistently monitored. The total duration of the whole imaging experiment was approximately 1.5 hours. For 1H MRS, adjustments of all first- and second-order shims within the voxel of interest were accomplished with the MAPSHIM procedure. The *in vivo* shimming procedure yielded a full width half maximum (FWHW) ranging from 7.8–10.9 Hz for the unsuppressed water peak in the spectroscopic voxel of the mouse brain. The water signal was attenuated by variable power radiofrequency (RF) pulses with optimized relaxation delays (VAPOR). Outer volume suppression (OVS) integrated with a point-resolved spectroscopy (PRESS) sequence was employed for signal acquisition, utilizing 2 x 2 x 2 mm^3^ voxel, with a TR/TE = 2500/20 ms, a spectral bandwidth of 4 kHz, 2048 data points, and 256 averages. Spectral data were acquired from the voxel encompassing the dorsal thalamic brain region including the PVT according to the mouse brain atlas. Metabolite quantification was conducted utilizing the LC-Model (version 6.3-1L) alongside the spectral basis set provided by the vendor. The absolute concentrations of the following metabolites were assessed: aspartate, creatine, γ-aminobutyric acid (GABA), glutamine, glutamate, guanidinoacetate, glycerophosphorylcholine, lactate, myo-inositol, N-acetylaspartate, N-acetylaspartylglutamate, Phosphocreatine, phosphorylcholine, and taurine. Only those results with Cramér-Rao lower bounds (CRLB, %SD) ≤ 50% were included in the statistical analysis.

### Repeated intermittent ethanol exposure

The repeated intermittent ethanol exposure paradigm has been described [37]. Mice were subjected to ethanol exposure in a vapor inhalation chamber [38] for three weeks, from postnatal day 28 to postnatal day 46. Each daily’s cycle included either 16 hours of ethanol vapor (AIE group, BEC 100-150 mg/dL at the end of the session) or equivalent counterpart room air (CON group), followed by 8 hours of abstinence in their home cage away from both vaporized ethanol and air. This was repeated daily for four consecutive days, succeeded by three days of abstinence. The cellular activity and animal behaviors were assessed four weeks post-AIE paradigm (Fig. 1a).

**Fig 1.**
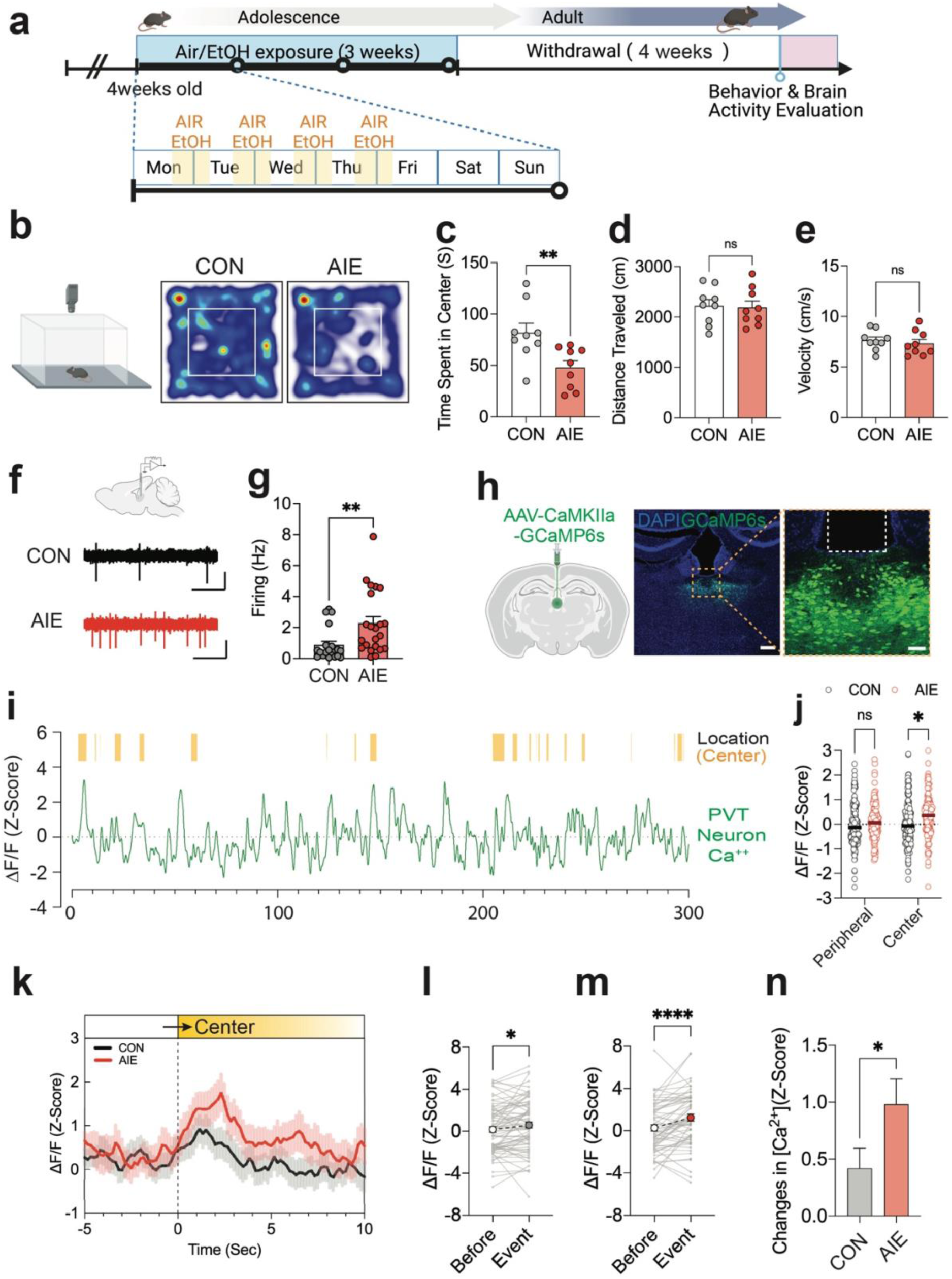
Adolescent repeated ethanol exposure (AIE) induces anxiety-like behaviors in adult mice and increased neuronal activities in the PVT. Diagram to explain the experimental paradigm (a). Representative traces (b) and pooled data (c-e) showing that adult mice at 4 weeks withdrawal from repeated ethanol exposure during adolescent period (AIE) show heightened anxiety-like behaviors compared to air-exposed counterpart mice (CON) (Fig. 1c, Unpaired t-test, CON vs AIE, t=2.966, df=16, p=0.0091; Fig. 1d, Unpaired t-test, CON vs AIE, t=0.1698 df=16, p=0.8673; Fig. 1e, Unpaired t-test, CON vs AIE, t=0.5992, df=16, p=0.5574, N_mice_=9/group). (f-g) Diagram and representative traces (f) and pooled data (g) showing that increased spontaneous firing of PVT neurons after AIE (Fig. 1g, Unpaired t-test, CON vs AIE, t=2.738, df=40, p=0.0092, N_cell_=20-22 [5 mice/group]). (h-n) Representative expression of GCaMP6s in the PVT (h), behavior-synced calcium traces (i), and pooled data (j-n) showing that the calcium transients in the PVT neurons are significantly increased in the AIE group (Fig. 1j, Two-way RM ANOVA, Interaction: F(1,643)=1.065, P=0.3026, Group: F(1,643)=5.629, P=0.0180, Location: F(1,643)=9.629, P=0.0020, Center:CON vs Center:AIE, p=0.0195, N_events_=126-197/group [4 mice/group]; Fig. 1l, Paired t-test, Before vs Event, t=2.381, df=73, p=0.0199, N_events_=74/group; Fig. 1m, Paired t-test, Before vs Event, t=4.413, df=61, p<0.0001, N_events_=62/group; Fig. 1n, Unpaired t-test, t=2.008, df=134, p=0.0466, N_events_=62-74/group [4 mice/group]). Data represented as mean ± SEM. *p<0.05, **p<0.01, ****p<0.0001.

### Behavioral Evaluations

All home cages housing mice were transported to the behavior testing room a minimum of 2 hours before the testing start. All the tests were scored using video tracking software, Ethovison (Ethovision XT, Noldus).

#### Open field test

The open-field test (OFT) was performed in a chamber (40 x 40 x 40 cm) to assess anxiety-like behaviors and locomotion. The session went for 30 minutes in the testing room under low light intensity (∼30 lux). The duration in the center area (25 x 25 cm), distance to travel, and velocity were compared as 5-minute time bins, with the initial 5-miute interval serving as a reference for assessing anxiety-like behavior.

#### Light-dark box test

The light-dark box test (LDT) was performed in a chamber designed as a modified open-field arena (60 x 40 x 30 cm), where about two-thirds of the apparatus is brightly lighted (∼550 lux) and the remaining one-third is devoid of light (∼3 lux). To create the different light intensity in the two areas, the illumination in the testing room was deactivated (0 lux), and LED lights were affixed to the wall separating the two areas, directed towards the brightly illuminated open area. The animal was placed in the non-illuminated section of the apparatus to freely explore both sections for 10 min; time spent in the lit section, distance traveled, and velocity were recorded as 5-minute time bins and the first 5-min bin was used to compare the anxiety levels between groups.

#### Elevated plus maze test

Elevated plus maze (EPM) was positioned 50 cm above the ground and comprised two open (35 cm x 6 cm x 0.5 cm) and two closed (35 cm x 6 cm x 22 cm) arms, together with a connecting central zone (6 x 6 cm). Mice were placed in the closed arm of the EPM, facing to the end of the arm. Mice were allowed to navigate the maze for 5 minutes, during which the duration spent in the open arms, distance traveled, and, velocity were recorded.

#### Tail suspension test

To evaluate depressive-like behaviors, mice were suspended by their tails with adhesive tape from a bar positioned 20 cm above the table. Individual mice were partitioned by a barrier, and the tests were conducted for a duration of 6 minutes. The total area moved, activity duration, and duration of time spent immobile during the last 4 minutes were analyzed. Mice were considered immobile when they hung passively without any movement.

#### Forced swim test

Each mouse was placed in a 2 Liter beaker filled with water (30 cm depth, temperature 24-25°C) and forced to swim 6 mininutes. The duration of immobility for the last 4 minutes was recorded. They were considered immobile when they stopped struggling, only moved slightly and occasionally to keep their nose above the water surface.

#### Spontaneous alteration behavior in Y-maze

Spontaneous alternation, an assessment of spatial working memory, was conducted in a Y-maze arena (each arm: 30 X 6 X 15 cm). Mice were placed in one arm of a Y-maze, oriented towards the wall, and allowed 10 minutes to explore all three arms. The alternation ratio was calculated by dividing the number of complete visitings (A to B to C) by the total number of arms entered, subtracting two.

### Western blotting

Tissue containing the PVT was punched out from coronal slices (1 mm thick) and homogenized in ice-cold RIPA lysis buffer (Thermo Fisher) containing a protease inhibitor cocktail (Sigma Aldrich). Equal amounts of protein extracts were denatured and subjected to SDS-PAGE using 4-12% Bis-Tris gels and transferred to PVDF membranes (Thermo Fisher). PVDF membranes were blocked with tris-buffered saline containing 0.05% Tween 20 and 5% (w/v) non-fat dried milk. These membranes were incubated with anti-GLT1 antibody (1:2000, Guinea pig, Sigma Aldrich), anti-GLAST antibody (1:2000, Rabbit, Alomone Labs), anti-GAPDH antibody (1:2000, Mouse, Sigma Aldrich), and anti-xCT antibody (1:1000, Rabbit, ABclonal) overnight at 4 °C. After washing with TBST, the membranes were incubated for 1 h with appropriate horseradish peroxidase-conjugated secondary antibodies for 1 h at room temperature. The proteins were visualized by ECL solution (Thermo Fisher) using the G:Box Chemiluminescent Imaging System (Syngene).

### Glutamate assay

For assessing the glutamate levels in the brain tissue including the PVT, a colorimetric glutamate assay kit was used (ab83389, Abcam). The optical density (OD) value of each well was measured using a Spectrophotometer (Multiskan FC Photometer, Thermo Fisher) at an absorption wavelength of 450nm for each assay.

### Data analysis

All data are represented as mean ± SEM using Prism 10.1 (GraphPad Software, San Diego, CA). The statistical significance was set at *p* < 0.05. Detailed statistical tests and data with exact *p* values are listed in Supplementary Table 1.

## Result

### The anxiety-like behaviors in adult mice exposed to ethanol repeatedly during adolescent period are accompanied with hyperactivity of PVT neurons

To characterize the animal behaviors and brain adaptation induced by the adolescent chronic intermittent alcohol (ethanol) exposure (AIE) in adulthood, we exposed the mice to ethanol or counterpart air (CON) based on chronic intermittent ethanol exposure paradigm when the animals were four weeks old. The paradigm persisted for three weeks and ended before the animals escaped the adolescent period. After four weeks withdrawal from the last ethanol exposure, we subsequently evaluated the anxiety-like behaviors through open filed test (OFT), light-dark box test (LDT), and elevated plus maze test (EPM) (Figure 1 and Supplementary Figure 1). While the total distance traveled and movement velocity were similar in both groups, the time spent in the center area in the OFT was significantly shorter in the AIE group compared to that in the CON group (Figure 1b-e). Likewise, AIE group spent less time than the CON group in the bright light illuminated section during the LDT and in the open-arms during the EPM test (Supplementary Figure 2). Compared to the CON group, the AIE group mice did not exhibit significant changes in short-term spatial working memory in the spontaneous alternation behavior test (SAB) and depressive-like behaviors in the forced swimming test (FST) and tail suspension test (TST) at the four weeks withdrawal from AIE paradigm (Supplementary Figure 3), indicating that repeated alcohol exposure during adolescent period increased the susceptibility to anxiety-like behaviors in adult.

Next, we examined whether the AIE paradigm induces adaptation of the neuronal activities in the paraventricular nucleus of thalamus (PVT), an area that has been considered a hub of arousal and anxiety network [24,25,39], via electrophysiology and temporal calcium imaging with fiber-photometry. In the *in vivo* electrophysiology, the basal rate of spontaneous spiking was significantly higher in the PVT neurons of AIE mice than that of CON mice (Figure 1f-g). To further clarify whether the AIE-induced anxiety-like behaviors are related to the neuronal activities in the PVT, we measured spatiotemporal cellular activities of the PVT neurons with GCaMP6s, a genetically encoded Ca^2+^ indicator, synched with the behaviors in the OFT (Figure 1h-n). The results showed that, while the mice didn’t show any group differences in the calcium transients when they stayed in the peripheral area, AIE mice had enhanced calcium transients in the center area compared to that of CON group (Figure 1i-j). In addition, AIE mice had much stronger calcium transients in the PVT neurons than that in the CON mice during the exploratory behaviors to the center area in the OFT (Figure 1k-n). These data suggest that the hyperactivity of PVT neurons is accompanied with AIE-induced anxiety-like behaviors.

### Chemogenetic inhibition of the PVT neurons alleviates the anxiety-like behaviors induced by AIE in adulthood

To determine whether the PVT neuronal adaptation contributes to the anxiogenic phenotype in adulthood of the AIE mice, we sought to check whether chemogenetic inhibition of the PVT neurons will ameliorate the AIE-induced anxiety-like behaviors. We infected the PVT neurons with an adeno-associated viral vector (AAV) expressing inhibitory DREADDs, hM4Di, under the control of a CaMKIIa promoter, leading to the selective expression of hM4Di in the projecting, excitatory PVT neurons [40–42] (Figure 2a-b). In *ex vivo* electrophysiological recordings, we confirmed the application of the DREADDs ligand, JHU31760 (J60, 20 μM), effectively silenced the firing of PVT neurons expressing hM4Di (Figure 2c). We then found that the administration of J60 (0.3 mg/kg, i.p.) significantly increased the time spent in the center in the hM4Di-expressed AIE mice, without the significant changes in total distance traveled or velocity during the open field test (Figure 3d-g), suggesting that PVT inhibition could alleviate the anxiety-like behaviors in adult induced by AIE.

**Fig 2.**
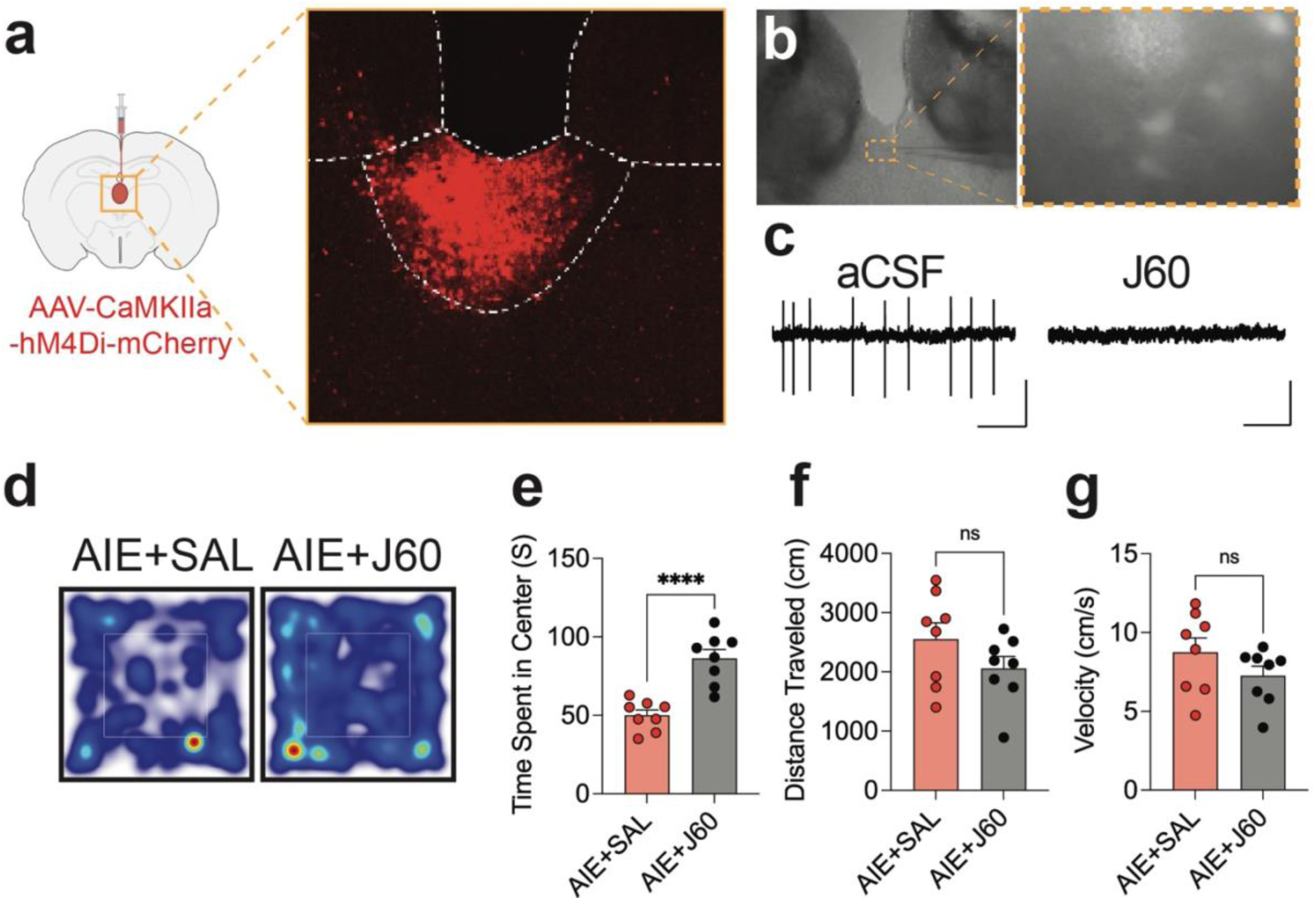
Chemogenetic inhibition of PVT neurons alleviates anxiety-like behaviors induced by AIE. (a-b) Representative figures showing the expression of hM4Di-mCherry in the PVT. (c) Representative traces showing the silence of neuronal activity in the hM4Di-positive neurons in the PVT by bath application of DREADDs ligand (J60, 20 µM). (d-g) Represenetative traces (d) and pooled data (e-g) showing that the inhibition of PVT neuronal activities by chemogenetic application rescues the anxiety-like behaviors seen in AIE mice (Fig. 2e, Unpaired t-test, AIE+SAL vs AIE+J60, t=5.511, df=14, p<0.0001, N_mice_=8/group; Fig. 2f, Unpaired t-test, AIE+SAL vs AIE+J60, t=1.448, df=14, p=0.1697, N_mice_=8/group; Fig. 2g, Unpaired t-test, AIE+SAL vs AIE+J60, t=1.359, df=14, p=0.1955, N_mice_=8/group). Data represented as mean ± SEM. ****p<0.0001. J60: JHU37160 (0.3 mg/kg, *i.p.*).

**Fig 3.**
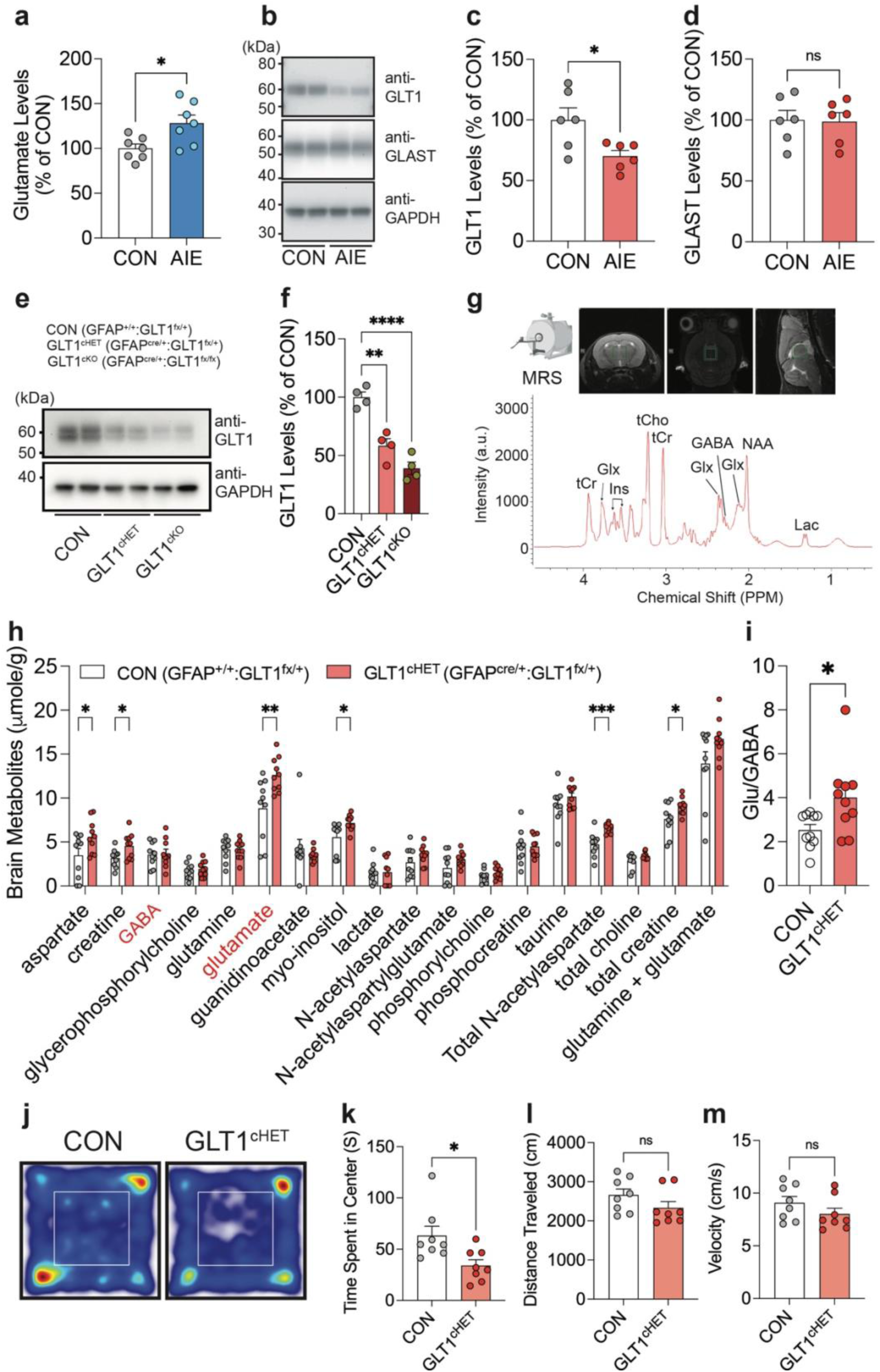
The expression of GLT1, an astrocytic glutamate transporter, is selectively reduced in AIE mice and GLT1 conditional knockdown induces anxiogenic phenotypes. (a) The quantification of glutamate levels in the dorsal thalamic area including PVT (Unpaired t-test, CON vs AIE, t=2.757, df=12, p=0.0174, N_mice_=7/group). (b-d) Representative expression of GLT1 (as known as EAAT2), GLAST (as known as EAAT1), and GAPDH (b) and pooled data (c-d) of western blots showing that the GLT1 expression in the PVT of AIE mice is selectively reduced compared to that of the CON mice (Fig. 3c, Unpaired t-test, CON vs AIE, t=2.725, df=10, p=0.0214, N_mice_=6/group; Fig. 3d, Unpaired t-test, CON vs AIE, t=0.1174, df=10, p= 0.9089, N_mice_=6/group). (e-f) Representative figures (e) and pooled data (f) confirming the reduction of GLT1 in the conditional GLT1 knockdown mice (GLT1^cHET^) (One-Way ANOVA, Treatment: F(2,9)=34.20, P<0.0001; Tukey’s posthoc, CON vs. GLT1^cHET^, p=0.001, CON vs. GLT1^cKO^, p<0.0001, GLT1^cHET^ vs. GLT1^cKO^, p=0.0716). (g-i) Non-invasive magnetic resonance spectroscopy (MRS) measurement showing the increase in glutamate levels in the dorsal thalamic area including PVT of GLT1^cHET^ (Fig. 3h, Unpaired t-test, CON vs GLT1^cHET^, aspartate: t=2.144, df=18, p=0.0460; creatine: t=2.476, df=18, p=0.0235; γ-aminobutyric acid [GABA]: t=0.1341, df=18, p=0.8948; glycerophosphorylcholine: t=0.5625, df=18, p=0.5807; glutamine: t=0.1383, df=18, p=0.8916; glutamate: t=3.006, df=18, p=0.0076; guanidinoacetate: t=0.7702, df=18, p=0.4512; myo-inositol: t=2.624, df=18, p=0.0172; lactate: t=0.08816, df=18, p=0.9308; N-acetylaspartate: t=1.403, df=18, p=0.1777; N-acetylaspartylglutamate: t=1.646, df=18, p=0.1171; Phosphorylcholine: t=1.173, df=18, p=0.2559; Phosphocreatine: t=0.1111, df=18, p=0.9128; Taurine: t=1.351, df=18, p=0.1934; Total N-acetylaspartate: t=4.017, df=18, p=0.0009; total choline: t=2.019, df=18, p=0.0587; total creatine: t=2.253, df=18, p=0.037; glutamine + glutamate: t=1.867, df=18, p=0.0784, N_mice_=10/group; Fig. 3i, CON vs GLT1cHET, t=2.497, df=18, p=0.0225, N_mice_=10/group). (j-m) Representative traces (j) and pooled data (k-m) showing the anxiogenic profiles of GLT1^cHET^ (Fig. 3k, Unpaired t-test, t=2.700, df=14, p=0.0173; Fig. 3l, Unpaired t-test, t=1.472, df=14, p=0.1632; Fig. 3m, Unpaired t-test, t=1.322, df=14, p=0.2073; N_mice_=8/group). Data represented as mean ± SEM. *p<0.05, **p<0.01, ***p<0.001, ****p<0.0001.

### Involvement of glutamatergic signaling between astrocytes and neurons in the AIE-induced adaptation of the PVT

Previous studies have demonstrated a hyper-glutamatergic state in brain, characterized by elevated extracellular glutamate, underlies increased susceptibility to chronic ethanol exposure and withdrawal-induced neurotoxicity and synaptic activity [43]. Specifically, astrocytic glutamate transporters, which have an eminent role to scavenge the extra glutamate in the tripartite synapses, have been identified as a factor of brain adaptation that induces anxiety-like behaviors [44,45] and those malfunctional adaptations are frequently observed after repeated ethanol exposure and withdrawal [46,47]. Thus, to determine whether the capacity of glutamatergic signaling in the PVT is changed by AIE, we quantified glutamate levels and the protein levels of GLT1 and GLAST, the two major astrocyte-dominant transporters, in the PVT. First, we found the PVT tissue of AIE mice had an increase in glutamate levels compared to those of CON (Figure 3a). Interestingly, this increase in glutamate levels was coupled with a significantly reduced expression in GLT1 in the PVT tissue of AIE mice compared to that of CON (Figure 3b and c), while the GLAST expression remained unchanged (Figure 3b and d), suggesting that AIE induced hyper-glutamate levels in the PVT, at least partly, via the adaptation of GLT1.

To assess how the GLT1 reduction affects metabolites in the brain area including glutamate *in vivo*, we measured the metabolites non-invasively using magnetic resonance spectroscopy (MRS) in the mice with a GFAP-driven conditional heterozygous (cHET) reduction of GLT1 in the astrocytes (GLT1^cHET^, GFAP^cre/+^:GLT1^fx/+^; Figure 3e-f and Supple figure 4). Conditional targeting was verified with GFAP-promoter driven GLT1 conditional knock-out mice (GLT1^cKO^, GFAPCre/+:GLT1fx/fx; Figure 3e-f). Heterozygous reduction in GLT1 is sufficient to attenuate the GLT1 levels without significant changes in GLAST and xCT levels in the PVT, similar to the changes by AIE (Figure 3e and f and Supplementary Figure 4). This heterozygous reduction in GLT1 also excludes the lethal phenotype shown in GLT1 cKO and global GLT1 KO [48,49]. The glutamate level in the thalamic area, including the PVT, of the mice was significantly reduced without changes in GABA levels, which induces the enhanced balance between glutamate and GABA ratio (Figure 3g-i). When we measured their behaviors in the OFT, the time spent in the center was significantly reduced in the GLT1^cHET^ mice compared to that in the CON mice, with no change in distance traveled or velocity (Figure 3j-m and Supplementary Figure 5).

**Fig 4.**
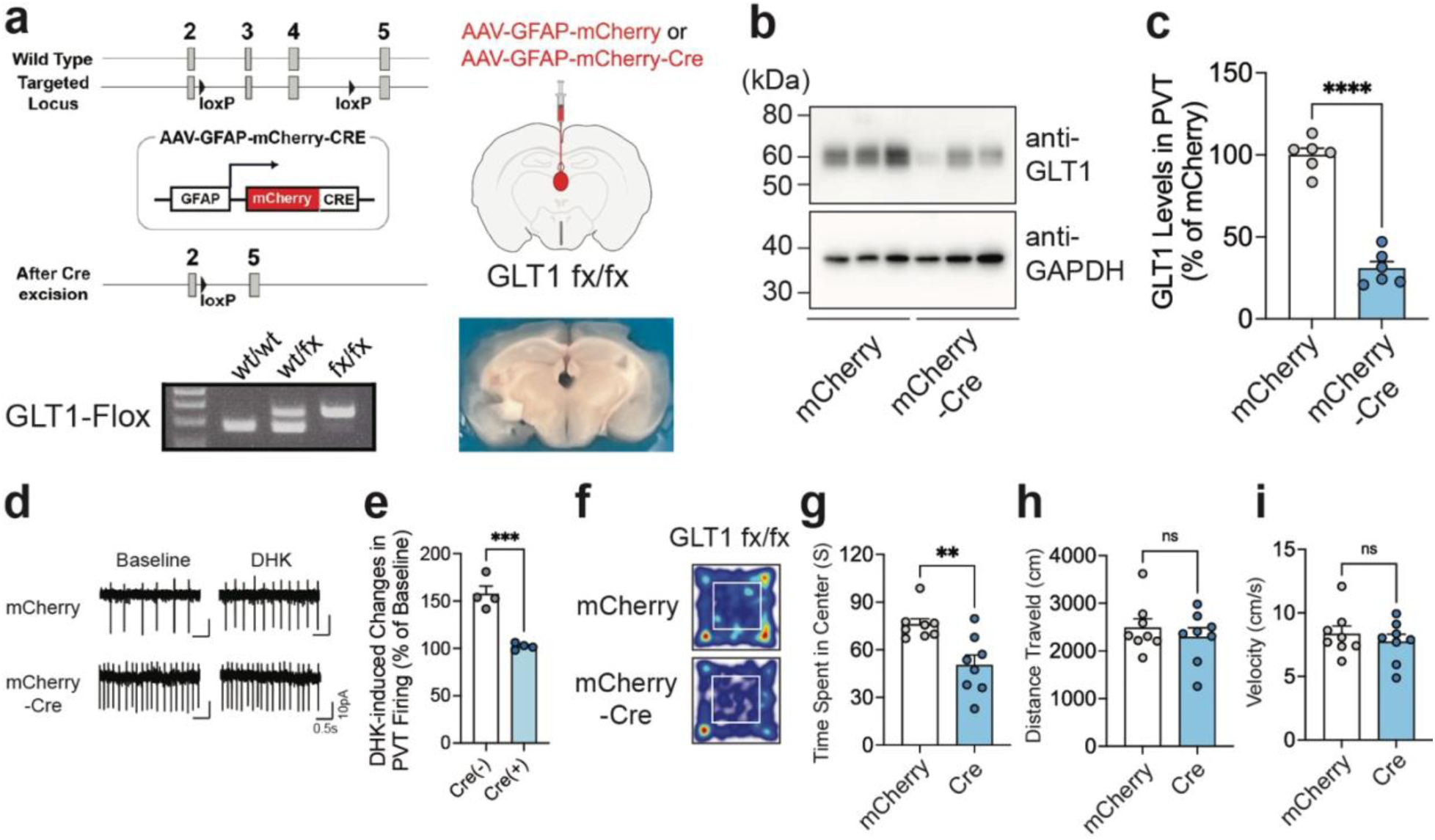
Region-specific conditional knock-out of GLT1 in the PVT mimics the anxiety-like behaviors induced by adolescent ethanol exposure. (a) Schematic drawing of Cre-loxP system for GLT1 cKO and genotyping of GLT1 wild-type and floxed mouse. (b,c) The representative figures (b) and pooled data (c) showing the reduction of GLT1 levels in the PVT after the GFAP-promoter driven expression of Cre in the PVT of GLT1 floxed mouse. (d-e) Representative traces (d) and pooled data (e) showing that the DHK-induced changes in neuronal firing were occluded in the PVT of GLT1 cKO (Fig. 4e, Unpaired t-test, Cre[−] vs Cre[+], Unpaired t-test, t=6.166, df=6, p=0.0008). (f-i) The pooled data showing the effects of GLT1 cKO in the PVT astrocytes in the open field test (Fig. 4g, Unpaired t-test, mCherry vs Cre, t=3.471, df=14, p=0.0037; Fig. 4h, Unpaired t-test, mCherry vs Cre, t=0.7261, df=14, p=0.4797; Fig. 4i, Unpaired t-test, mCherry vs Cre, t=0.7156, df=14, p=0.486; N_mice_=8/group). Data represented as mean ± SEM. **p<0.01, ****p<0.0001.

Given the role of GLT1 reduction in anxiogenic phenotype, we sought to determine whether site-specific conditional knock-out of GLT1 in PVT induces the anxiogenic behavioral phenotype. We injected an AAV that has a capacity of a GFAP-promoter driven Cre recombinase expression in astrocytes into the PVT of ethanol naïve GLT1 floxed mice (Figure 4a and Supplementary Figure 6). The mice with the Cre recombinase expression showed a significant reduction in GLT1 in the astrocytes of the PVT compared to mCherry controls (Figure 4b and c). To further determine the functional adaptation of GLT1 cKO in the PVT, we compared the effects of Dihydrokainic acd (DHK), a selective GLT1 inhibitor in the PVT neurons of CON and AIE mice (Figure 4d-f). Bath application of DHK (200 μM) significantly increased the neuronal spiking of the PVT neurons in the CON. However, the changes by the bath-applied DHK in the PVT neurons were significantly reduced in the AIE. In the open field test, the mice that astrocyte GLT1 in the PVT was deleted showed the decrease in the time spent in the center in the open filed test without significant changes in locomotion (Figure4 f-i), suggesting that selective knock-out of GLT1 in the PVT astrocytes induces the anxiogenic phenotype.

Collectively, these pharmacological and transgenic validations suggest that the observed increases in glutamate levels and consequent behavioral changes could result from the lower expression and function of GLT1 in the PVT after AIE experience.

### Selective Rescue of GLT1 expression in the PVT of AIE mice ameliorate the AIE-induced anxiety-like behaviors

Because the AIE reduced the GLT1 expression in the PVT and induced the hyper-glutamate levels, which was accompanied with anxiogenic phonotypes, we sought to determine whether restoring GLT1 expression in the PVT would rescue the behavioral adaptation in AIE. We injected an AAV to express Cre recombinase in astrocytes using the GFAP promoter into the PVT of the AIE experienced transgenic Ai9-GLT1 mice (Figure 5a). The Ai9-GLT1 mice have a capacity to delete the stop-codon Cre recombinase dependently, leading a continuous increase in GLT1 expression (Figure 5a-d). In the open field test, the mice overexpressing astrocyte GLT1 became anxiolytic compared to mCherry-expressed AIE mice (Figure 5e-h and Supplementary Figure 7), indicating that GLT1 overexpression in the PVT of AIE mice ameliorated the anxiogenic phenotype as observed in AIE mice.

**Fig 5.**
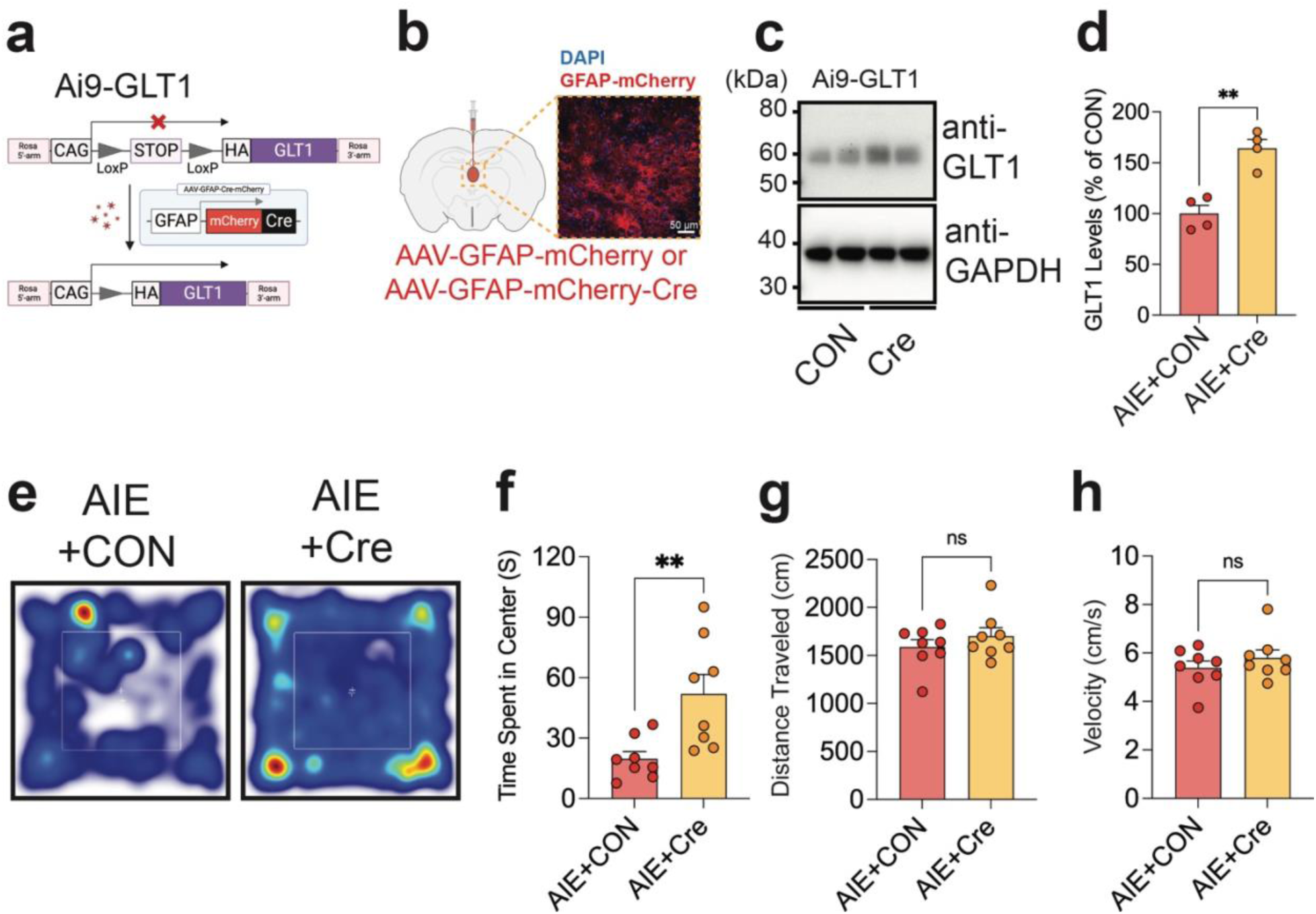
Rescue of GLT1 expression in the astrocytes of the PVT ameliorates the AIE-induced anxiety-like behavior. (a) Transgenic approach to overexpress GLT1 in the astrocytes of PVT in a cell-type specific manner. (b-d) Diagram and representative expression (b), representative blots (c), and pooled data (d) showing the increased GLT1 expression in the PVT after the AAV-induced Cre expression in the PVT of the Ai9-GLT1 AIE mice (Fig. 5d, Unpaired t-test, AIE+CON vs AIE+Cre, t=5.398, df=6, p=0.0017, N_mice_=4/group). (e-f) Representative traces €, and pooled data (f-h) showing that the astrocytic GLT1 rescue in the PVT ameliorates the AIE-induced anxiety-like behavior in the open field test compared to the counterpart AIE mice (Fig. 5f, Unpaired t-test, AIE+CON vs AIE+Cre, t=3.143, df=14, p=0.0072; Fig. 5g, Unpaired t-test, AIE+CON vs AIE+Cre, t=0.9703, df=14, p=0.3484; Fig. 5h, Unpaired t-test, AIE+CON vs AIE+Cre, t=0.9500, df=14, p=0.3582; N_mice_=8/group).

Taken together, our findings support the hypothesis that adolescent repeated ethanol exposure alters the glutamatergic communication between astrocytes and neurons in the PVT, leading to the anxiogenic phenotypes in adult, at least partly, via the astrocyte glutamate transporter, GLT1.

## Discussion

In the present study, we provide comprehensive evidence of apparent anxiety-like behaviors in adult mice that were exposed ethanol repeatedly in their adolescent period. Importantly, chemogenetic inhibition of the PVT neurons or transgenic overexpression of GLT1 in the PVT ameliorate the anxiety-like behaviors. In addition, while the PVT neuronal activity and glutamatergic signals are enhanced, the GLT1 expression is selectively reduced in the PVT. When GLT1 levels in the PVT are upregulated, the anxiogenic phenotype in adulthood that was induced by adolescent repeated ethanol exposure were rescued. These finds suggest that GLT1 in the PVT plays a significant role in the susceptibility of anxiety-related behaviors after the withdrawal from adolescent repeated ethanol exposure.

Similar to the previous reports of the anxiogenic effects of ethanol on rodents [16,17,50–53], we described anxiety-like behaviors in mice withdrawn from repeated ethanol exposure during the adolescent period, as shown by the reduced time spent in the center of an open field, and in the open arms of an elevated plus maze, as well as the less time in the light illuminated area. The total distance traveled remained unchanged, indicating the adolescent repeated ethanol exposure did not significantly impact the locomotor activity in our model. Notably, we observed the adaptation of cellular activities in the PVT, which is a main upstream brain region of central amygdala, where has been considered a hub of anxiety-network and reported to have brain adaptation induced by long-term withdrawal from adolescent intermittent ethanol exposure [53].

Although the PVT has been receiving increasing attention to be linked with anxiety-related behaviors, especially behavioral malfunction induced by adverse experiences during early-in life [20], its role in the context of withdrawal from adolescent repeated ethanol exposure was unknown. We found that PVT neurons in ethanol-withdrawn adults (AIE) had a significantly higher spontaneous firing rates, suggesting a possible contribution of PVT hyperactivity to the increased anxiety levels. This possibility was supported by our data showing that selective inhibition of PVT neurons by chemogenetic approach or transgenic upregulation of GLT1 in the PVT mitigated elevated anxiety-like phenotypes associated with the ethanol withdrawal. Searching for the underlying cellular and molecular mechanisms, we found that the PVT neurons of AIE mice had an enhanced level of glutamate and glutamate/GABA ratio, compared to CON mice and this is accompanied with the reduction of GLT1 levels in the PVT, suggesting that GLT1 dysregulation may contribute to the observed increased activities of PVT neurons in the AIE mice. These data support a link between PVT activity and anxiety-like behaviors.

The observation of prolonged brain adaptation including the reduction of GLT1 protein expression in AIE mice may be caused by changes in the factors that affect the corresponding gene expression. For example, the transcription factor, Pax6, has been identified as a common mechanism of regulating GLT1 expression, without expression on the levels of another astroglial glutamate transporter, GLAST, by binding within the upstream of the GLT1 translation start site [54]. Indeed, the Pax6 is expressed early in development, predominantly in a few body parts including the brain [55], and has an essential role for early animal development [56,57]. While the expression of Pax6 is reduced after ethanol exposure [58], Pax6 overexpression rescued alcohol-induced cellular dysfunction [59]. In addition, multiple epigenetic mechanisms via microRNAs, which are short non-coding RNAs modulating mRNA translation, have been suggested to contribute to the brain adaptation due to alcohol exposure during adolescence, which increase the susceptibility in adult to alcohol use and anxiety [50,53,60–62]. Thus, given the role of miRNAs and epigenetic impacts in the regulation of transcription factors, including Pax6 [63–65], it is intriguing to investigate the mechanisms how the synergistic reciprocal epigenetic modulations with related transcription factors affect the selective changes in GLT1 of the PVT by the adolescent repeated ethanol.

Although we did not identify the cell type we recorded in the current study, it has been known that the neurons in the PVT are majorly glutamatergic [66–68]. Recently, accumulative observations have provided the additional information there are diverse neuronal subpopulations in the PVT according to significant transcriptomic variance corresponding to the anterior-posterior axis [69,70], which may contribute to difference muti-brain regional connectivity and diverse behaviors. Importantly, PVT also has been well known to have a role of pivotal region for anxiety-network and have direct connections to multiple regions within the anxiety-network including the Central Amygdala (CeA) and Nucleus Accumbens (NAc) [25], where are the largest cluster of anxiety-network in the rodent and primate brains, participating in mediation of generalized anxiety and related arousal behaviors [71,72], and environment-raised anxiety and reward motivation behaviors [73,74], respectively. However, the functional significance of the various sub-connections from the PVT during ethanol withdrawal, specially adapted by the repeated exposure to ethanol during adolescent period, remains poorly addressed. Thus, future studies thus are needed to clarify which cell-types and circuits are more significantly impacted by adolescent repeated ethanol exposure in an age-dependent manner. In addition, given the role of PVT in arousal and vigilance-linked behaviors and the connection between anxiety and hyperarousal, it will be intriguing to investigate the role of PVT GLT1 in arousal-related behavioral patterns including social behaviors and sleep-wake cycles. One limitation of our study is the exclusion of sex differences by the use of both male and female mice. AIE induced behavioral changes are occasionally only observed in one sex [13]. Thus, age- and sex-dependent evaluation need to be further studied as well.

Our data here suggests that GLT1 downregulation-induced PVT hyperactivity in adults exposed to adolescent repeated ethanol exposure may contribute to anxiety-like behaviors. This study implies the importance of glutamatergic homeostatic balance through GLT1 in the adult behavioral susceptibility induced by the early life adverse experience.

## Data availability

All data are available from the authors upon reasonable request.

## Acknowledgment

We especially thank to Drs. Doo-Sup Choi and Niels Danbolt for providing GLT1-flox mouse and Drs. Doo-Sup Choi and Ho Lee for providing Ai9-GLT1 mouse. We also thank to Drs. Karl Deisseroth and Bryan Roth for providing plasmid DNA. We thank to Dr. Mungyu Song, Kishan Trivedi, Vasu Kapoor, Sahil Patel, Lesley Barksdale, Ni Na Ngo, Kaleb Louisy, Bansari Patel, and Chloe Yuri Woo for technical assistance and critical discussion. We express appreciation to all laboratory members for their valuable discussions and comments. Some of the figures were created with BioRender.com. This research was supported by the National Institute of Health (AA027773, MH137204 to SK).

## Author contributions

SK and AB designed the study. SK, AB, and HK performed all behavioral, electrophysiological, and biochemical experiments. SK and AB performed the stereotaxic surgeries. SK, AB, PB, and HK collected, processed, and imaged tissue for histology. RA, and AB performed and analyzed MRS. AB, DT, and SK wrote the manuscript. All authors reviewed and edited the manuscript.

## Supplementary Information

**Supplementary Fig. 1.**
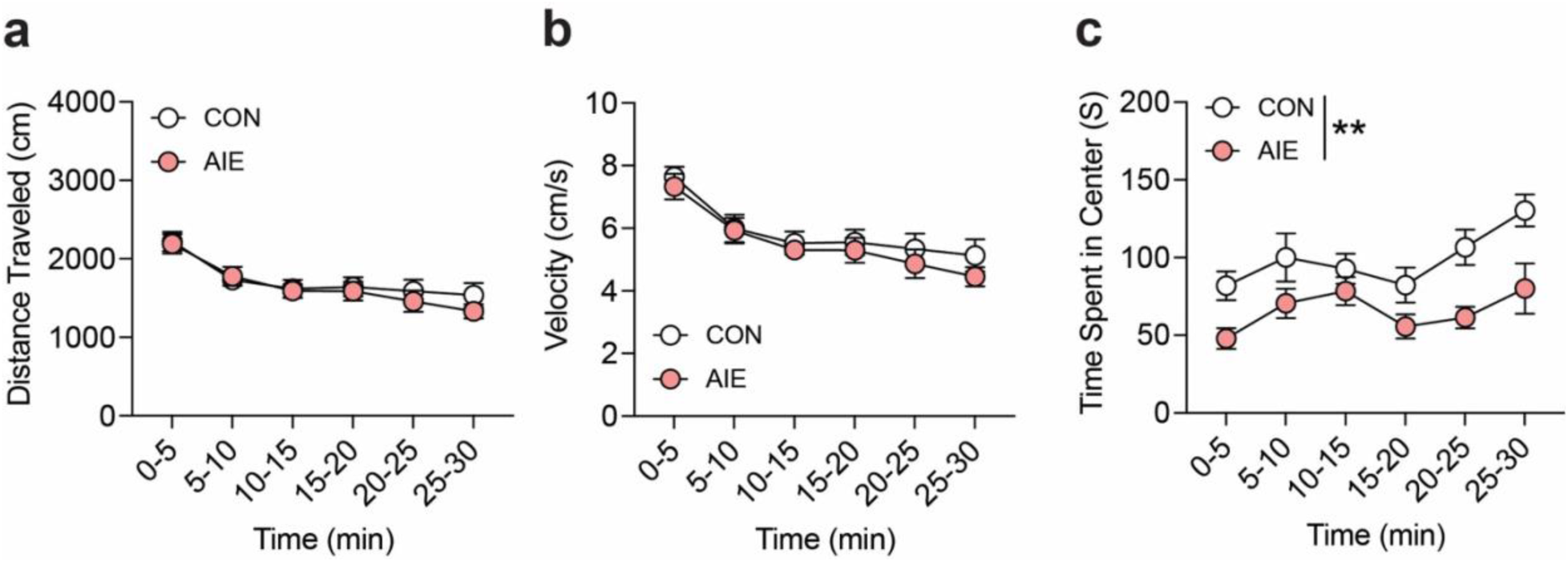
Adolescent repeated ethanol exposure (AIE) induces anxiety-like behaviors in adult mice. (a-c) pooled data showing the distance traveled (a), velocity (b), and the time spent in the center area (c) for total 30 minutes in the open-field arena. (Supplementary Fig. 1a, Two-way RM ANOVA, Time x Group: F (5, 80) = 0.4869, P=0.7851, Time: F (3.433, 54.92) = 18.66, P<0.0001, Group: F (1, 16) = 0.2691, P=0.6110, Subject: F (16, 80) = 6.131, P<0.0001, N_mice_=9/group; Supplementary Fig. 1b, Two-way RM ANOVA, Time x Group: F (5, 80) = 0.2514, P=0.9380, Time: F (3.339, 53.42) = 20.22, P<0.0001, Group: F (1, 16) = 0.6350, P=0.4372, Subject: F (16, 80) = 5.796, P<0.0001, N_mice_=9/group; Supplementary Fig. 1c, Two-way RM ANOVA, Time x Group: F (5, 80) = 1.160, P=0.3360, Time: F (3.918, 62.69) = 5.676, P=0.0006, Group: F (1, 16) = 10.43, P=0.0052, Subject: F (16, 80) = 4.400, P<0.0001, N_mice_=9/group). Data represented as mean ± SEM. **p<0.01.

**Supplementary Fig. 2.**
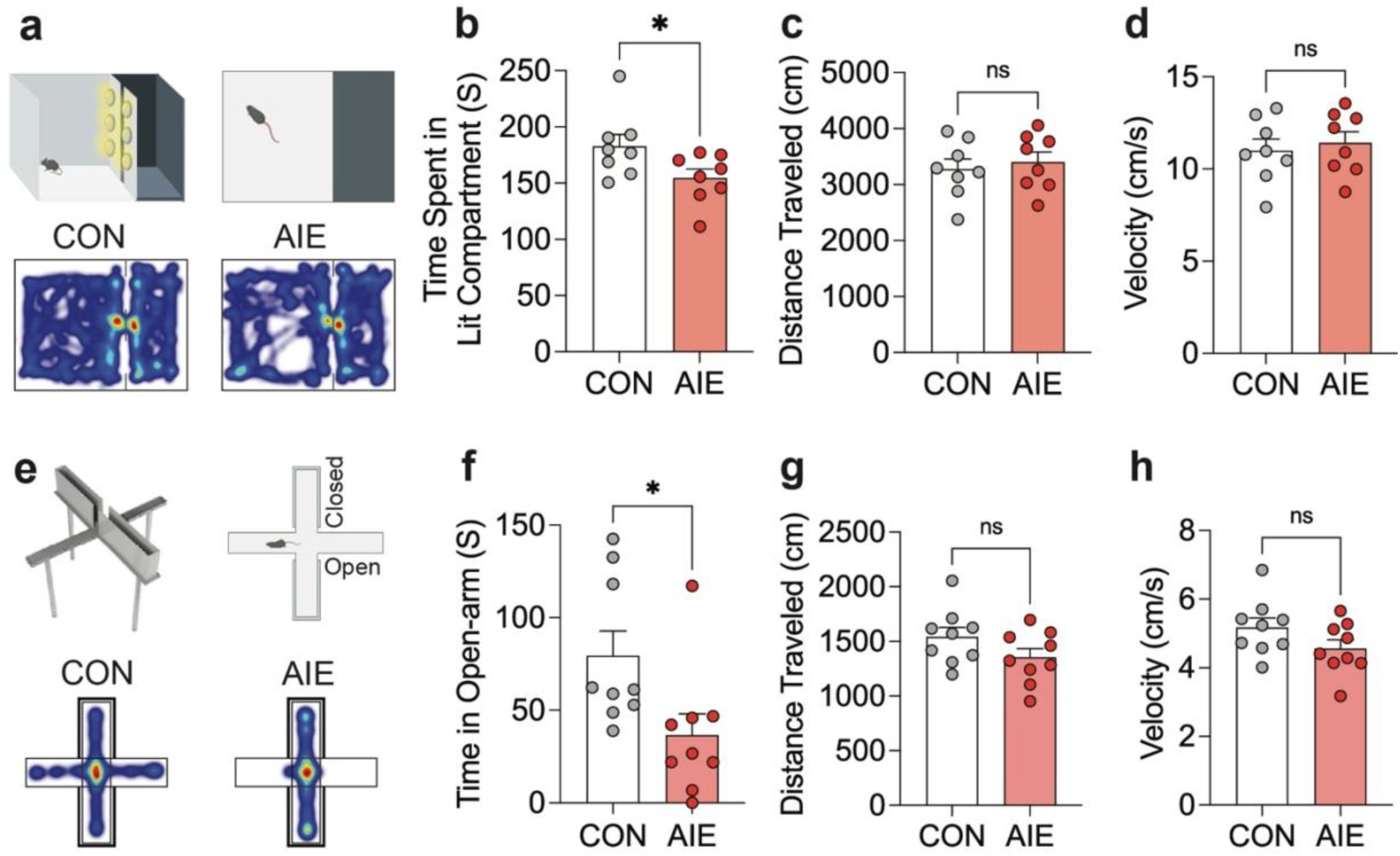
Adolescent repeated ethanol exposure (AIE) induces anxiety-like behaviors in adult mice. Representative traces and pooled data in the Light-dark box test (LDT, a-d, Supplementary Fig. 2b, Unpaired t-test, CON vs AIE, t=2.173, df=14, p=0.0474; Supplementary Fig. 2c, Unpaired t-test, CON vs AIE, t=0.4966, df=14, p= 0.6272; Supplementary Fig. 2d Unpaired t-test, CON vs AIE, t=0.4833, df=14, p= 0.6364, N_mice_=8/group) and elevated plus maze test (EPM, e-h, Supplementary Fig. 2f, Unpaired t-test, CON vs AIE, t=2.454, df=16, p= 0.026; Supplementary Fig. 2g, Unpaired t-test, CON vs AIE, t=1.616, df=16, p= 0.1257; Supplementary Fig. 2h, Unpaired t-test, CON vs AIE, t=1.664, df=16, p= 0.1157, N_mice_=9/group) showing that adult mice at 4 weeks withdrawal from repeated ethanol exposure during adolescent period (AIE) show heightened anxiety-like behaviors compared to air-exposed counterpart mice (CON). Data represented as mean ± SEM. *p<0.05.

**Supplementary Fig. 3.**
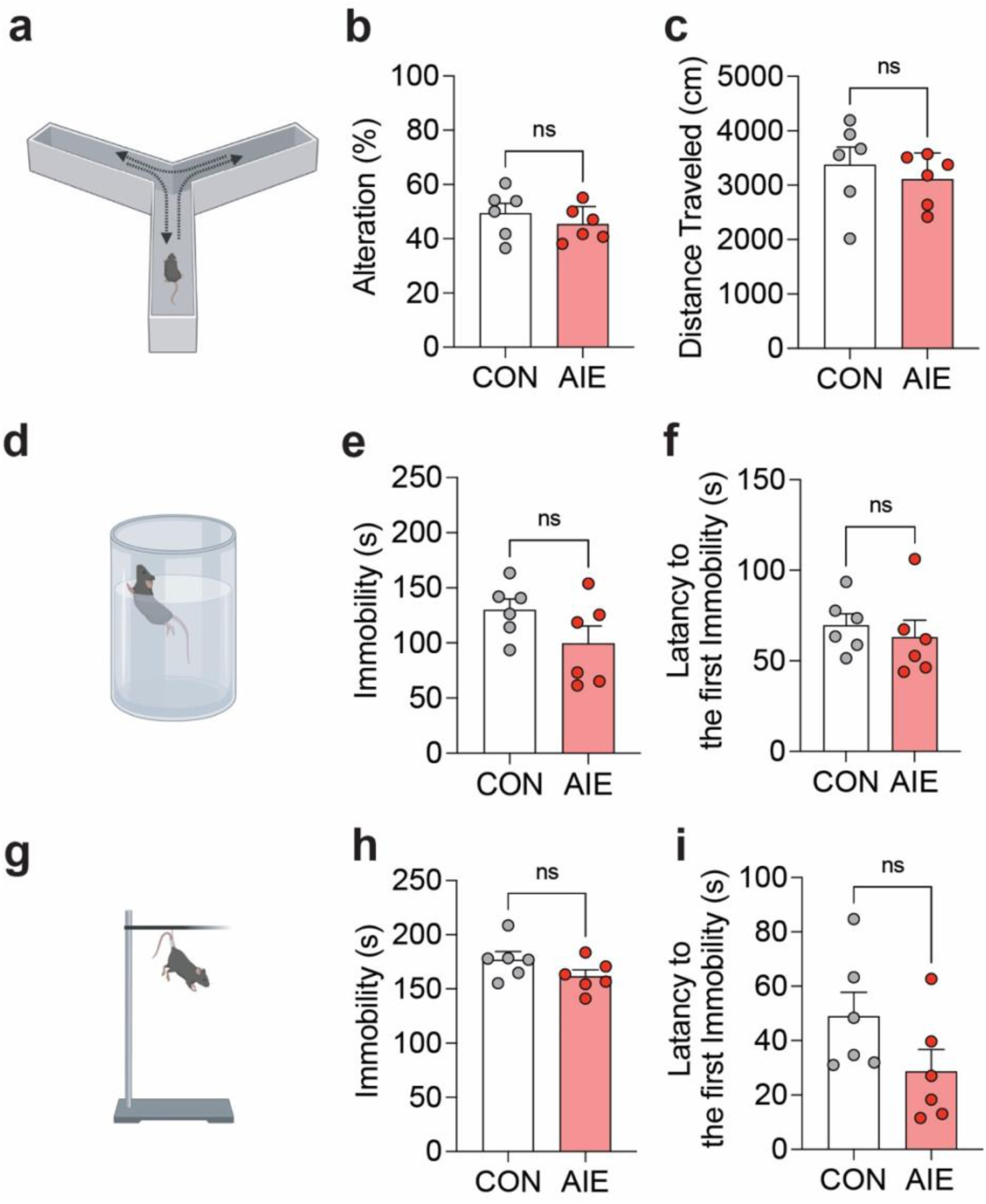
Adolescent repeated ethanol exposure (AIE) does not significantly affect working memory or induce depressive-like behaviors in adult mice. Representative traces and pooled data in the spontaneous alteration behavior in Y-maze (a-c, Supplementary Fig. 3b, Unpaired t-test, CON vs AIE, t=0.9017, df=10, p=0.3884; Supplementary Fig. 3c, Unpaired t-test, CON vs AIE, t=0.6829, df=10, p= 0.5102, N_mice_= 6/group), forced swim test (FST, d-f, Supplementary Fig. 3e, Unpaired t-test, CON vs AIE, t=1.643, df=10, p=0.1314, Supplementary Fig. 3f, Unpaired t-test, CON vs AIE, t=0.5990, df=10, p=0.5625, N_mice_=6/group), and Tail suspension test (TST, g-I, Supplementary Fig. 3h, Unpaired t-test, CON vs AIE, t=1.529, df=10, p=0.1572, Supplementary Fig. 3i, Unpaired t-test, CON vs AIE, t=1.710, df=10, p=0.1181, N_mice_=6/Group). Data represented as mean ± SEM.

**Supplementary Fig. 4.**
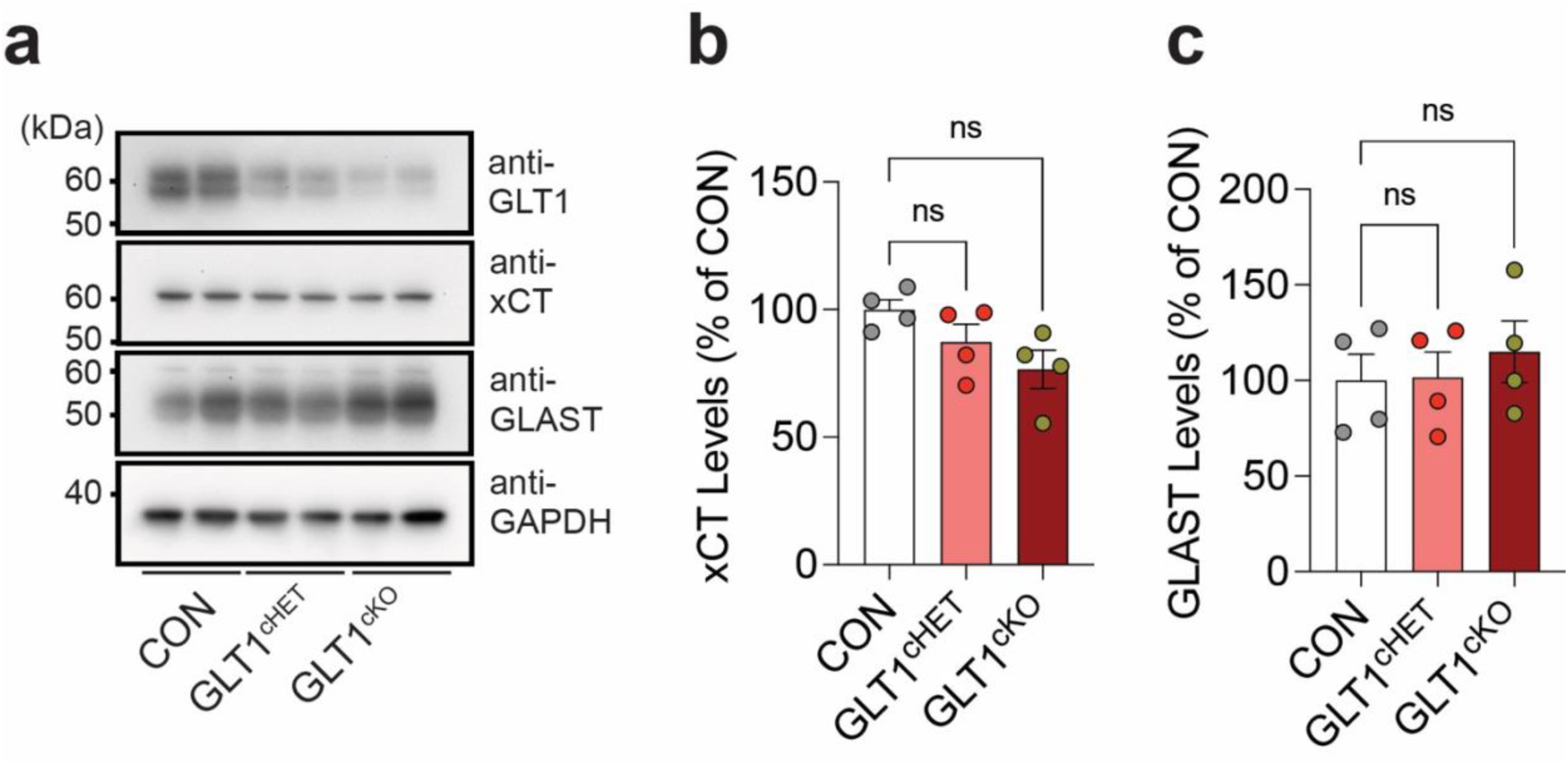
GLT1 conditional knockdown does not induce significant changes in xCT or GLAST levels. Representative blots (a) and pooled data (b-c) showing the levels of xCT (Supplementary Fig. 4b, One-way ANOVA, F (2, 9) = 3.492, P=0.0754, Tukey’s posthoc: CON vs. GLT1^cHET^, p=0.3683; CON vs. GLT1^cK^O, p=0.0634; GLT1^cHET^ vs. GLT1^cKO^, p=0.4756, N_mice_=4/group), and GLAST (Supplementary Fig. 4c, One-way ANOVA, F (2, 9) = 0.3250, P=0.7307, Tukey’s posthoc: CON vs. GLT1^cHET^, p=0.9962; CON vs. GLT1^cKO^, p=0.7492; GLT1^cHET^ vs. GLT1^cKO^, p=0.7954, N_mice_=4/group) in the PVT. Data represented as mean ± SEM.

**Supplementary Fig. 5.**
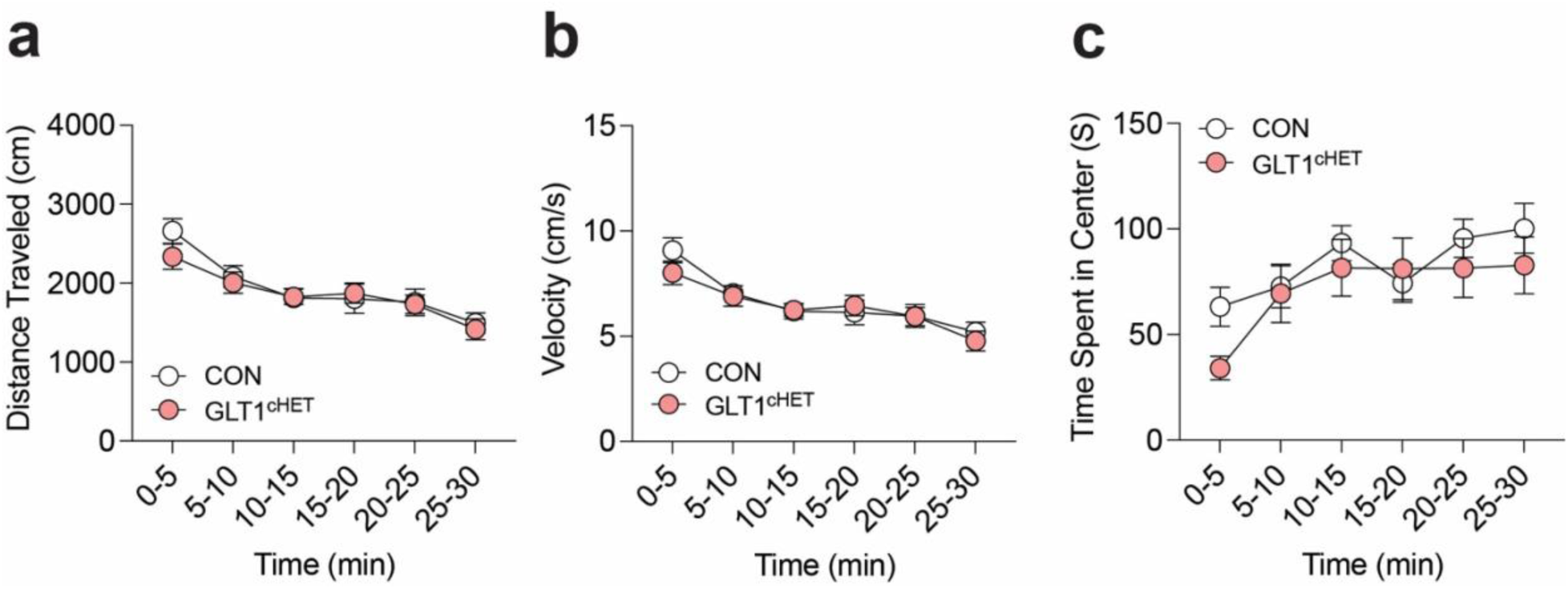
GLT1 conditional knockdown induces anxiogenic phenotypes. (a-c) pooled data showing the distance traveled (a, Supplementary Fig. 5a, Two-way RM ANOVA, CON vs GLT1^cHET^, Time x Group: F (5, 70) = 0.9783, P=0.4372, Time: F (3.181, 44.54) = 25.38, P<0.0001, Group: F (1, 14) = 0.2170, P=0.6485, Subject: F (14, 70) = 7.272, P<0.0001, N_mice_=8/group), velocity (b, Supplementary Fig. 5b, Two-way RM ANOVA, CON vs GLT1^cHET^, Time x Group: F (5, 70) = 1.073, P=0.3826, Time: F (3.334, 46.68) = 25.56, P<0.0001, Group: F (1, 14) = 0.1639, P=0.6917, Subject: F (14, 70) = 7.460, P<0.0001, N_mice_=8/group), and the time spent in the center area (c, Supplementary Fig. 5c, Two-way RM ANOVA, CON vs GLT1^cHET^, Time x Group: F (5, 70) = 1.028, P=0.4080, Time: F (3.431, 48.03) = 7.029, P=0.0003, Group: F (1, 14) = 0.9741, P=0.3404, Subject: F (14, 70) = 5.466, P<0.0001, N_mice_=8/group) for total 30 minutes in the open-field arena. Data represented as mean ± SEM.

**Supplementary Fig. 6.**
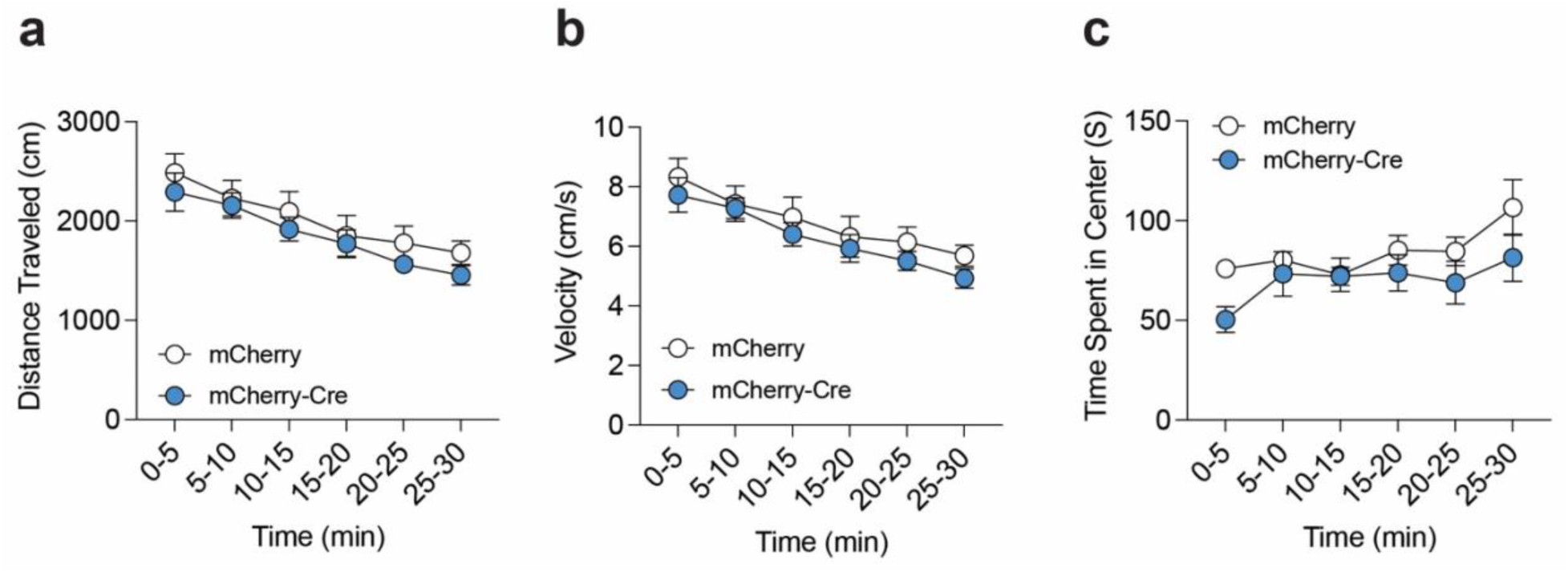
Region-specific conditional knock-out of GLT1 in the PVT mimics the anxiety-like behaviors induced by adolescent ethanol exposure (AIE). (a-c) pooled data showing the distance traveled (a, Supplementary Fig. 6a, Two-way RM ANOVA, mCherry vs mCherry-Cre, Time x Column Factor: F (5, 70) = 0.2531, P=0.9369, Time: F (3.534, 49.47) = 21.91, P<0.0001, Column Factor: F (1, 14) = 0.7697, P=0.3951, Subject: F (14, 70) = 11.18, P<0.0001, N_mice_=8/group), velocity (b, Supplementary Fig. 6b, Two-way RM ANOVA, mCherry vs mCherry-Cre, Time x Column Factor: F (5, 70) = 0.2323, P=0.9471, Time: F (3.323, 46.52) = 20.48, P<0.0001, Column Factor: F (1, 14) = 0.7780, P=0.3926, Subject: F (14, 70) = 10.47, P<0.0001, N_mice_=8/group), and the time spent in the center area (c, Supplementary Fig. 6c, Two-way RM ANOVA, mCherry vs mCherry-Cre, Time x Column Factor: F (5, 70) = 0.7429, P=0.5940, Time: F (3.015, 42.21) = 3.072, P=0.0377, Column Factor: F (1, 14) = 4.644, P=0.0491, Subject: F (14, 70) = 1.980, P=0.0320, N_mice_=8/group) for total 30 minutes in the open-field arena. Data represented as mean ± SEM.

**Supplementary Fig. 7.**
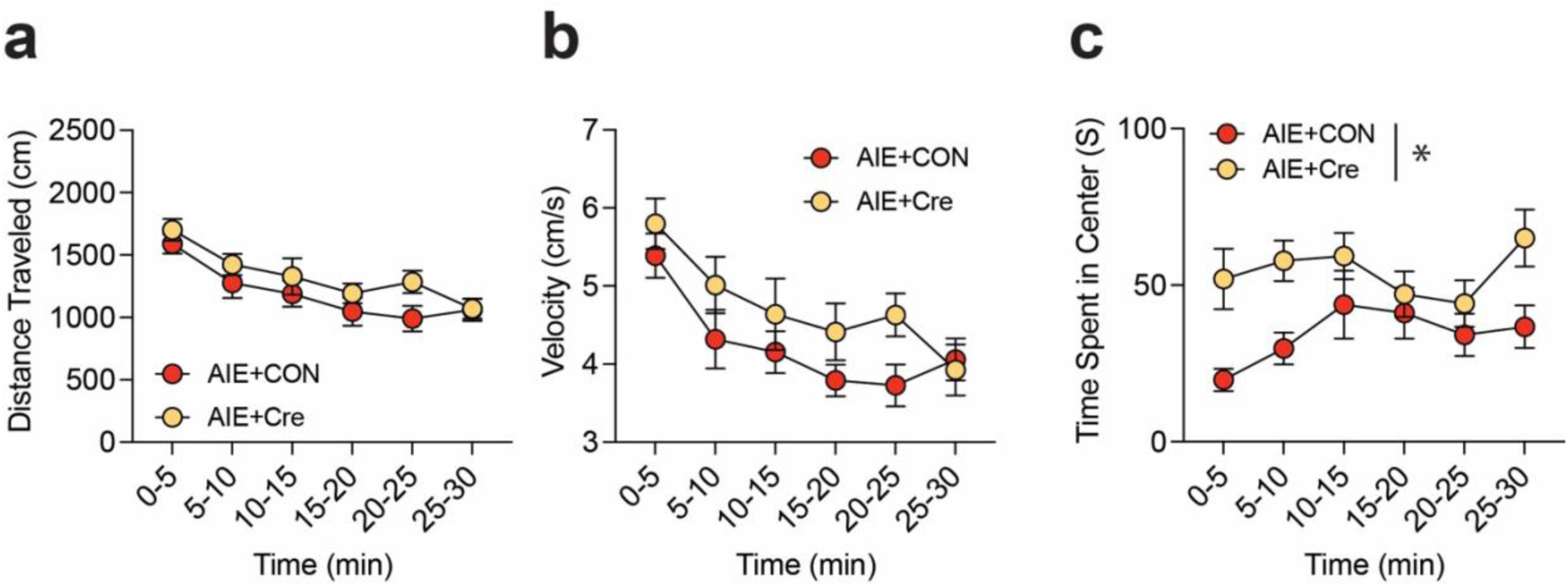
Rescue of GLT1 expression in the astrocytes of the PVT ameliorates the AIE-induced anxiety-like behavior. (a-c) pooled data showing the distance traveled (a, Supplementary Fig. 7a, Two-way RM ANOVA, AIE+CON vs AIE+Cre, Time x Group: F (5, 70) = 0.7829, P=0.5654, Time: F (3.613, 50.58) = 17.09, P<0.0001, Group: F (1, 14) = 1.843, P=0.1960, Subject: F (14, 70) = 6.100, P<0.0001, N_mice_=8/group), velocity (b, Supplementary Fig. 7b, Two-way RM ANOVA, AIE+CON vs AIE+Cre, Time x Group: F (5, 70) = 0.9670, P=0.4441, Time: F (3.389, 47.44) = 10.84, P<0.0001, Group: F (1, 14) = 2.506, P=0.1357, Subject: F (14, 70) = 4.559, P<0.0001, N_mice_=8/group), and the time spent in the center area (c, Supplementary Fig. 7c, Two-way RM ANOVA, AIE+CON vs AIE+Cre, Time x Group: F (5, 70) = 1.721, P=0.1410, Time: F (2.744, 38.42) = 2.235, P=0.1046, Group: F (1, 14) = 6.931, P=0.0197, Subject: F (14, 70) = 4.981, P<0.0001, N_mice_=8/group) for total 30 minutes in the open-field arena. Data represented as mean ± SEM. *p<0.05.

